# Whole-brain connectomics of Drosophila reveals a robust, distributed architecture for the suppression of feeding during escape

**DOI:** 10.64898/2025.12.14.694122

**Authors:** Wei-Qiang Chen, Wenjie Xi

**Affiliations:** Department of Physics and HK Institute of Quantum Science & Technology, The University of Hong Kong, Pokfulam Road, Hong Kong, China; Department of Physics and Shenzhen Key Laboratory of Advanced Quantum Functional Materials and Devices, Southern University of Science and Technology, Shenzhen 518055, China

## Abstract

Survival demands instant prioritization of escape over maintenance. To decode the dynamic logic embedded in the static Drosophila connectome (FlyWire v783), we simulated the conflict between predator-evasion, feeding, and grooming to uncover the logic of this switch. We show that escape is a holistic state defined by distributed robustness: it is reliably triggered by only a fraction of visual inputs (LC4/LPLC2) and induces a system-wide pause on behaviors like feeding and grooming. Within this global suppression, we identify a specialized, fail-safe architecture for arresting feeding. A redundant ensemble of neurons (DNge031/CB0565) targets the premotor center Roundup, effectively disfacilitating motor output. Crucially, this inhibition is modular, anatomically distinct from grooming suppression, and relies on additive logic to ensure suppression even if individual components fail. Our results demonstrate how connectome topology implements robust, survival-critical control through distributed neural architecture.

## Introduction

To survive in a complex environment, animals must continuously prioritize mutually exclusive behaviors, executing high-priority actions such as predator evasion while suppressing maintenance drives such as feeding [1, 2]. A central tenet of neuroethology attributes this rapid switching to “command neurons”—single critical nodes whose activation is both necessary and sufficient to trigger a specific behavioral program [3–5]. This reductionist paradigm spans diverse species. For example, the Giant Fiber (GF) system in Drosophila [6, 7], the Mauthner cells in fish [4], and lateral giant neurons in crayfish [8] are all classically defined as dedicated decision nodes for escape. Similarly, in mammals, specific hypothalamic populations such as AgRP neurons are often modeled as centralized switches that drive feeding [9]. However, this focus on single-node control presents an engineering paradox. Biological systems display remarkable resilience to noise and damage [10]. Such holistic stability is incompatible with architectures reliant on a single critical node [11, 12]. Theoretical work on biological complexity suggests that robust-ness arises not from singular controls but from degeneracy—the ability of structurally distinct elements to perform the same function [13]—and distributed population codes [14]. Despite these theoretical insights, the specific synaptic architectures that implement such distributed robustness within the brain remain largely unmapped.

The recent completion of the adult Drosophila whole-brain connectome (FlyWire) [15, 16] provides an unprecedented opportunity to resolve this paradox. By mapping every synaptic connection, we can now identify distributed networks that were previously invisible to sparse sampling methods. Furthermore, a static wiring diagram alone cannot predict which pathways are functionally recruited during specific behaviors. This data set allows us to investigate the holistic topology of neural states. Crucially, this perspective reveals that a behavior is rarely executed by a solitary command. For instance, a successful escape requires not just the trigger (e.g., DNp01 [6, 17]), but the co-activation of postural control (e.g., DNp04 [7]) and directional steering (e.g., DNa02 [18, 19]) to form a complete motor synergy. To decode the dynamic logic embedded in this static map, we apply a connectome-constrained spiking neural network (SNN) model of the entire brain [20]. This full-brain simulation approach enables us to perform “virtual stress tests” on the network—systematically perturbing inputs and performing combinatorial silencing of hidden layers to reveal the redundancy and stability of behavioral states. This allows us to dissect fault-tolerant architectures that are difficult to probe via single-gene manipulations in vivo. This framework allows us to ask not just how a behavior is triggered, but how the system ensures its execution is robust against internal variability and external perturbations.

In this paper, we simulate the competition between predator evasion and maintenance circuits including feeding and grooming to study the circuit logic of high-priority behavioral switching. We reveal that the escape response is not a rigid reflex driven by isolated lines, but a holistic neural state protected by distributed stability at both input and output levels. First, we show that escape is reliably initiated by activating only a fraction of the looming-sensitive population (LC4/LPLC2) [7,21,22], demonstrating that danger detection relies on robust population coding rather than critical individual inputs. Second, we identify a specialized, fail-safe architecture that enforces behavioral exclusivity. We find that the escape circuit suppresses feeding not through lateral inhibition at the motor output (MN9), but via upstream disfacilitation. A redundant ensemble of descending (DNge031) and central (CB0565) neurons collectively inhibits Roundup, the primary upstream of NM9, effectively cutting off the feeding drive at its source. This architecture offers a distinct control advantage: by suppressing the premotor drive rather than competing at the muscle level, the system avoids metabolic waste and prevents “jittery” motor conflicts, ensuring a clean state transition. By removing the excitatory input from Roundup rather than inhibiting the motor neuron MN9 directly, this mechanism prevents conflicting signals from competing at the output stage, ensuring a efficient behavioral switch. Perturbation analysis reveals that this inhibition operates on an additive logic: the failure of any single inhibitor is insufficient to release the feeding drive, establishing a redundant safety margin that guarantees the absolute priority of survival. Finally, we demonstrate that this inhibitory architecture is highly modular. While escape induces a system-wide pause, it utilizes distinct mechanisms for different behaviors: the disfacilitation of Roundup is specific to feeding, whereas grooming is suppressed via separate, dedicated gating pathways. Our findings demonstrate that the connectome is engineered for systemic robustness, where stable behavioral switching emerges from the synergistic action of distributed neural assemblies.

## Results

In this paper, we applied a full-scale SNN model [20] constrained by the adult Drosophila brain connectome (FlyWire v783) [15,16] to study the circuit logic of behavioral switching. This model incorporates 127,400 proofread neurons and approximately 50 million synaptic connections, representing the complete wiring diagram of the adult Drosophila brain. The dynamics of the model is characterized by a Leaky Integrate-and-Fire (LIF) formalism, a standard approach that balances computational tractability with biophysical realism. The model was constrained strictly based on anatomical data: synaptic weights were proportional to the number of ultrastructural synapses between neuron pairs, and the sign of the interaction (excitatory or inhibitory) was determined by brain-wide neurotransmitter predictions [23]. By driving the model with stochastic Poisson inputs, we transcend static graph analysis to simulate the dynamic transformation of sensory signals into coordinated motor outputs.

### Holistic Activation of Escape

we simulated a “looming predator” by activating the visual projection neurons LC4 and LPLC2. Biological studies have established that these neurons act as specific feature detectors, summing their inputs to drive the GF membrane potential [7]. Accordingly, we drove these inputs at high frequencies (150 Hz) to mimic the peak synaptic drive generated during the terminal phase of a rapid loom. Biologically, escape is not a singular muscle twitch but a coordinated motor synergy involving takeoff triggering, postural adjustment, and directional steering. Our simulation accurately captured this holistic activation pattern across varying input frequencies (Fig1a).

**Figure 1.**
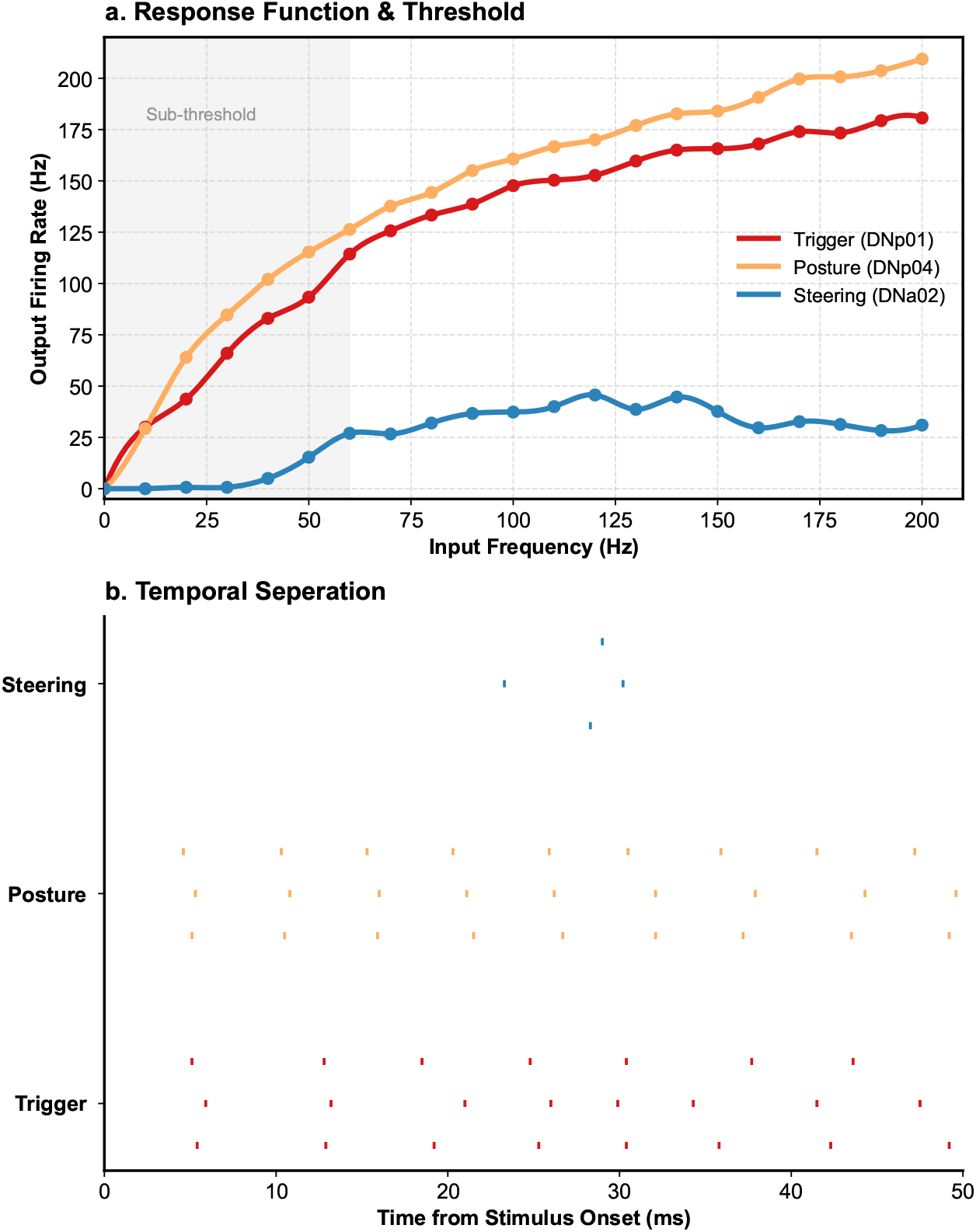
Frequency-response curve and temporal dynamics of the escape synergy. (a) Frequency-response curves for the escape components: Trigger (DNp01, red), Posture (DNp04, orange) and Steering (DNa02, blue), generated with input activation fixed at 70% for both LC4 and LPLC2. The shaded region indicates firing rates rise sharply to a regimes (>60 Hz), demonstrating a non-linear response that ensures the escape synergy is stably elicited only under high-urgency conditions. (b) Spike raster (inputs= 150HZ) plot showing the temporal response of key escape components to a simulated high-urgency looming stimulus (start at *t* = 0). High-resolution analysis reveals a precise **temporal separation**: Trigger and Posture modules fire in tight synchronization with an ultra-short latency (∼5 ms) to initiate takeoff, while the Steering module exhibits a distinct delay (activating at ∼22 ms). This sequence suggests a motor logic that prioritizes takeoff before engaging directional flight control.

In response, the model GF (DNp01) reliably entered a high-frequency firing state (> 150Hz). While the biological GF typically triggers escape via a single action potential [7,17], its synapses are known to possess a high safety factor capable of following high-frequency stimulation [6]. Within our LIF framework, this sustained output represents a robust suprathreshold state, signifying a maximal excitatory drive that guarantees the “one-for-one” transmission required for instantaneous takeoff.

Crucially, we observed that the downstream activity was not confined to this primary line. The model spontaneously elicited a parallel ensemble of descending neurons, including DNp04(> 100 Hz) and DNp02(> 100 Hz), which are essential for postural stabilization and flight initiation [7]. High-resolution analysis reveals a precise temporal separation within this synergy (Fig1b). The trigger (DNp01) and postural (DNp04) modules fire in tight synchronization with an ultra-short latency (∼5 ms) to initiate takeoff. In contrast, the steering module (DNa02) exhibits a distinct delay, activating at∼22 ms. This temporal separation suggests that the connectome encodes a multi-stage motor program that prioritizes immediate lift-off before engaging directional flight control, optimizing the aerodynamics of escape. As shown in the co-activation function (Fig.2a), these distinct functional modules exhibit a synchronized onset, confirming that the connectome encodes a tightly coupled motor synergy. Furthermore, the steering command exhibited absolute directionality(Fig.2d): while right-sided visual input elicited robust activation of the contralateral (left) DNa02(> 20 Hz), the ipsilateral (right) DNa02 remained completely silent (0 Hz). This noise-free asymmetry provides an unambiguous, vector-correct signal for the fly to steer away from the threat, demonstrating that the connectome’s topology inherently encodes precise spatial computations.

**Figure 2.**
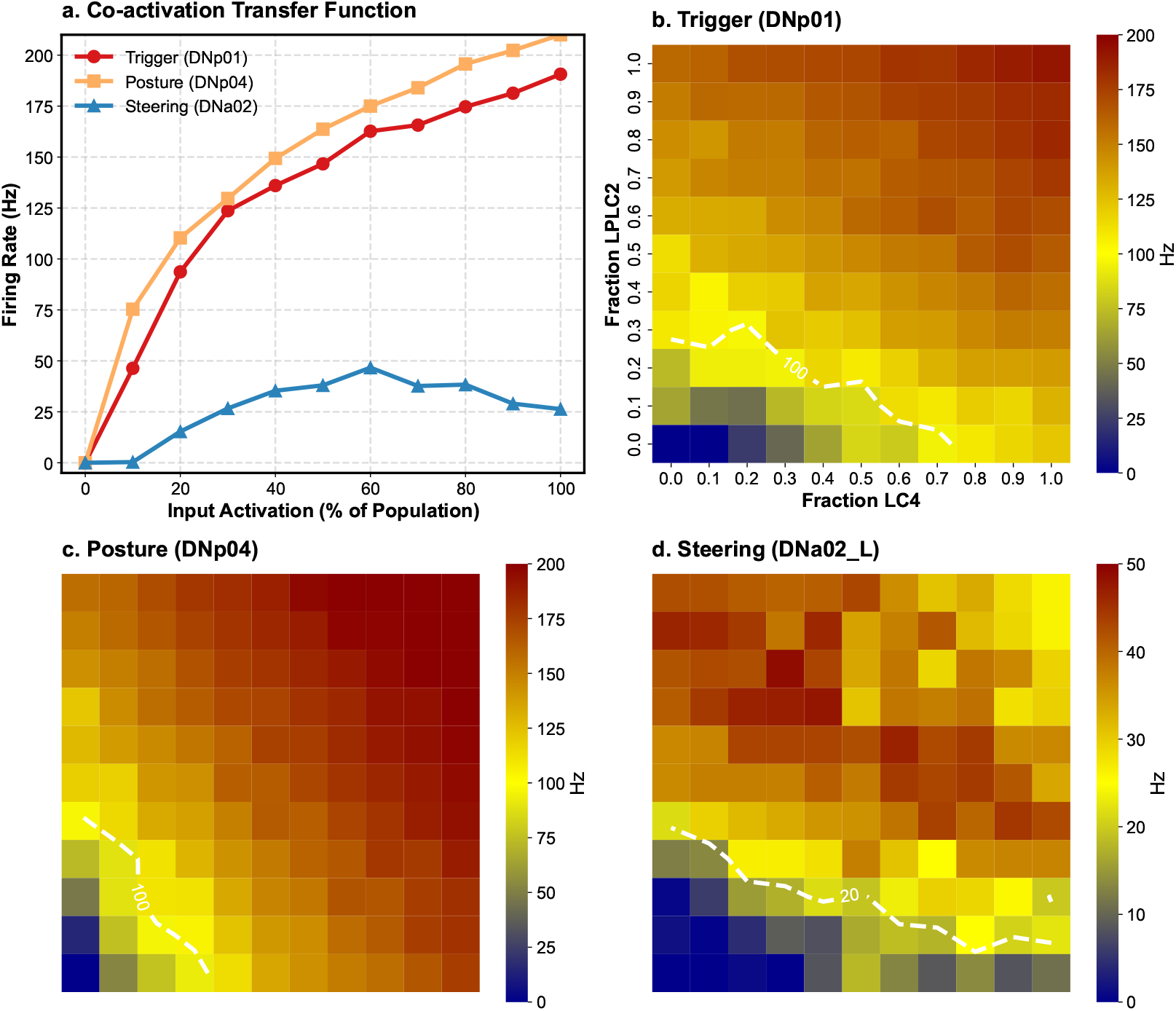
The synergistic and robust escape response. (a) Co-activation function showing the firing rates of three distinct functional modules: Trigger (DNp01, red), Posture (DNp04, orange), and Steering (DNa02, blue). These distinct motor programs are topologically locked into a unified behavioral state. (b-d) Input robustness landscapes (heatmaps) for each com-ponent. The color scale represents firing frequency (blue=silent, red=maximal). (b, c) The Trigger (DNp01) and Postural (DNp04) neurons exhibit a broad high-activity plateau (scale 0–200 Hz). The white dashed lines mark the 100 Hz threshold, demonstrating a wide safety margin where the behavior is reliably executed even if significant fractions of the input population (LC4/LPLC2) are silent. (d) The Steering neuron (left DNa02) shows a similar robust topology but operates in a lower frequency regime (scale 0–50 Hz, threshold 20 Hz), consistent with its role in providing continuous directional bias the right DNa02 is keeping silent.

### Robust Population Coding for escape

To determine the minimal input requirements for reliable escape initiation, we performed systematic “virtual stress tests” simulations by activating random fractions of the looming-sensitive population.

We observed a highly non-linear response profile where activating as few as 30% of LC4 and LPLC2 inputs was sufficient to drive the system into a stable “escape state” (Fig.2a). As visualized in the robustness landscapes, both Trigger (DNp01) and Postural (DNp04) neurons rapidly entered a broad high-activity plateau, maintaining firing rates well above the 100 Hz threshold even across varying input combinations (Fig.2b, c). Similarly, the Steering module (left DNa02) exhibited a robust directional bias, consistently firing above its 20 Hz threshold (Fig.2d) while the ipsilateral counterpart remained silent. This broad safety margin demonstrates that danger detection relies on distributed population coding: the escape response is not contingent on specific individual neurons but emerges from the collective activity of the visual assembly, ensuring robustness against noise and partial input loss. However, this potent and easily triggered escape state poses a regulatory challenge: how does the system prevent this high-gain signal from conflicting with simultaneous maintenance behaviors?

### Feeding suppression during escape

We next simulated a “conflict” scenario where the Drosophila is exposed to both sugar and a quickly looming threat. We found that triggering the escape response causes a rapid shutdown of feeding. As escape inputs (LC4/LPLC2) increase, the feeding motor neuron MN9 drops from high activity (> 80 Hz) to near-silence (Fig.3a). This creates a strict “exclusion zone” (dark blue region) where feeding is blocked. This establishes definitive evidence for the priority of survival, demonstrating that escape acts as a global state that forcibly overrides maintenance behaviors.

**Figure 3.**
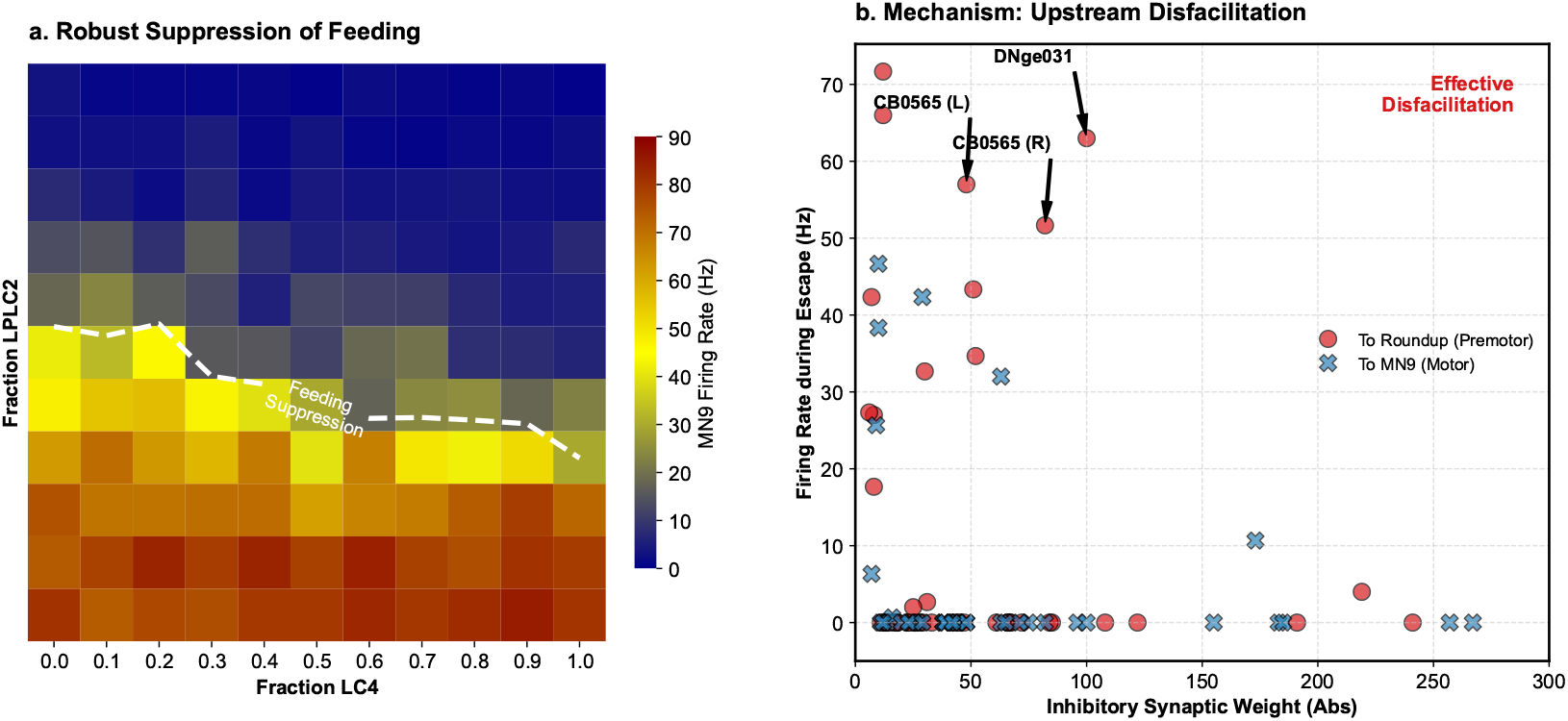
Escape suppresses feeding via targeted disfacilitation of the premotor center. (a) Heatmap showing the firing rate of the feeding motor neuron (MN9) during conflict (sugar stimulus + varying escape inputs). Despite constant sugar stimulation inputs (150 HZ), MN9 activity drops from high levels (> 80 Hz, red) to near-silence (blue) as escape inputs (LC4/LPLC2) increase. The white dashed line marks the suppression threshold (MN9< 30 Hz), showing a clear boundary where feeding stops. (b) Mechanism analysis for the escape state generated with input activation fixed at 100% for both LC4 and LPLC2 (150 HZ). Scatter plot comparing synaptic weight and firing rate for inhibitory neurons targeting either the motor neuron (MN9, blue crosses) or the premotor center (Roundup, red circles). The escape circuit selectively recruits strong inhibitors with high firing rates(e.g., DNge031, CB0565) targeting Roundup. In contrast, inhibitors targeting MN9 are either weak or inactive. This confirms that suppression is achieved by cutting off the excitatory drive at Roundup (upstream disfacilitation), rather than by inhibiting the motor neuron directly.

To find the mechanism behind this suppression, we traced the inhibitory signals of the escape circuit. Anatomically, the escape circuit connects to both the motor neuron (MN9) and its primary premotor driver, Roundup. However, effective suppression requires active neurons, not just static connections. By plotting synaptic weight against firing rate (Fig.3b), we revealed a clear functional difference. Inhibitors projecting directly to MN9 (blue crosses) were largely inactive or weak though they possessed high synaptic weights. In contrast, a specific group targeting Roundup, including the descending neuron DNge031 and central neuron CB0565, showed both high firing rates and strong inhibitory weights (marked red circles). This active inhibition silences Roundup, effectively cutting off the excitatory signals to MN9. This analysis exposes a critical limitation of purely static connectomics: anatomical connectivity does not equal functional inhibition. Therefore, the escape circuit enforces behavioral exclusivity via upstream disfacilitation, rather than directly inhibiting the motor neuron.

### Inhibitory architecture is highly robust and modular

To quantify the robustness of this suppression, we performed “counterfactual” simulations by silencing the identified inhibitors (DNge031 and CB0565) individually and in combination. We found that the system follows a strict additive logic (Fig.4b). Silencing any single inhibitor failed to restore feeding. Even silencing two out of three inhibitors resulted in only partial recovery (∼ 40 Hz), which remained significantly below the sugar-driven baseline (∼ 86 Hz). This safety margin ensures fail-safe termination of feeding, as the failure of any single component is insufficient to break the blockade.

**Figure 4.**
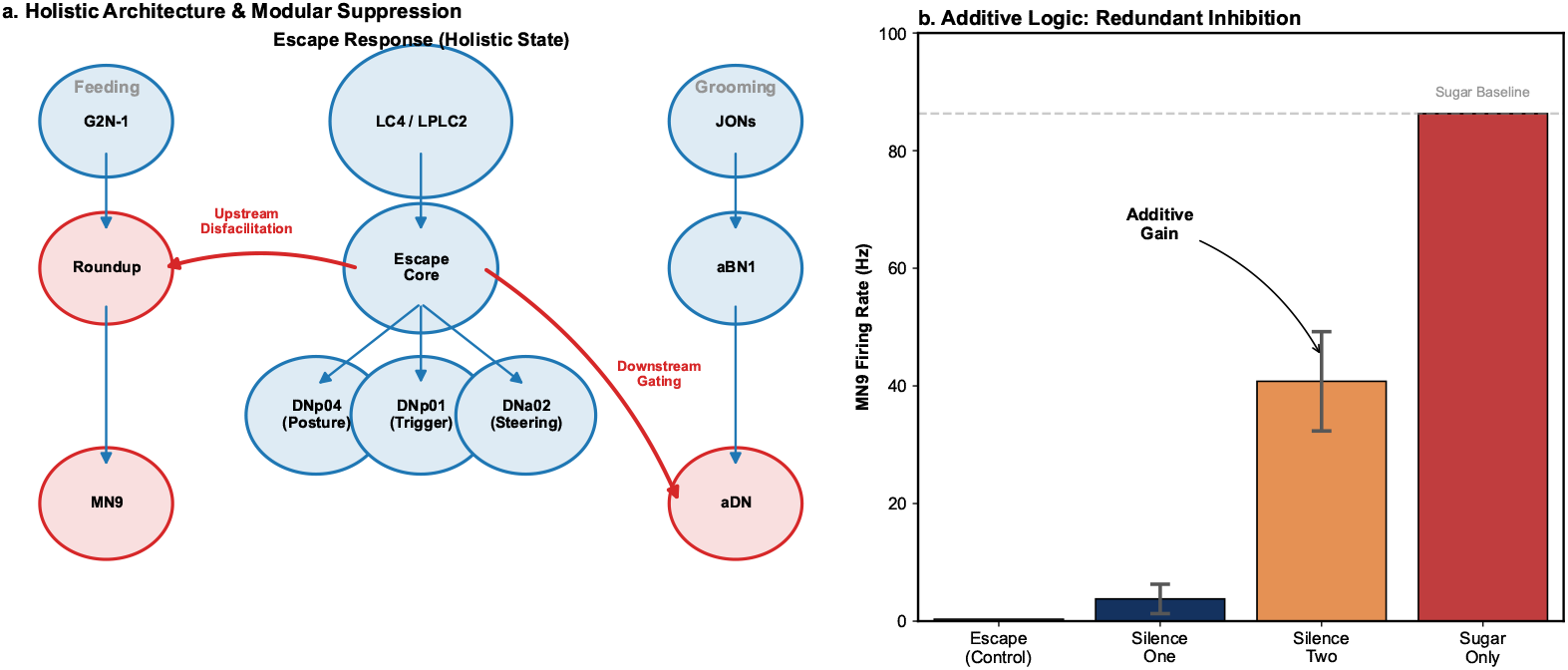
Modular specificity and additive redundancy of escape suppression. (a) Schematic of the holistic escape architecture where blue nodes indicate active neurons and red nodes indicate inactive/suppressed neurons. The escape core drives a synergistic motor output (center) while simultaneously suppressing competing behaviors via distinct, target-specific strategies. Feeding is suppressed via upstream disfacilitation (left): the escape core inhibits the premotor center Roundup (red), cutting off the excitatory drive to MN9. In contrast, grooming [24,25] is suppressed via downstream gating (right): the premotor drive (aBN1, blue) remains active, but the motor output (aDN, red) is blocked. (b) MN9 firing rates under combinatorial silencing of the identified inhibitors (DNge031, CB0565s) with input activation fixed at 100% for both LC4 and LPLC2 (150 HZ). Silencing any single inhibitor (“Silence One”) fails to restore feeding. Silencing two inhibitors (“Silence Two”) results in only partial recovery, demonstrating an additive gain. This redundancy creates a safety margin, ensuring fail-safe suppression even if individual inhibitory components fail.

Finally, we asked whether this architecture represents a global “freeze” or a targeted control policy. By comparing the feeding circuit with the grooming circuit, we found distinct, modular strategies (Fig.4a). While escape pauses both behaviors, it targets them at different hierarchical levels. Feeding is cut off at the source via upstream disfacilitation of Roundup. In contrast, the grooming premotor center (aBN1) remains active during escape, but its downstream output (aDN) is blocked via downstream gating. This functional dissociation confirms that the escape state acts through modular control policies, applying specific stopping mechanisms adapted to different behavioral conflicts.

## Discussion

### From Static Maps to Dynamic Logic

The FlyWire connectome provides a static map of the brain’s hardware. However, understanding behavioral logic requires dynamics. By applying a biophysical LIF model to this structure, we successfully recapitulated emergent properties, such as the temporal separation between ballistic takeoff and directional steering, which are not explicitly visible in the wiring diagram alone. This validates that connectome-constrained modeling is a sufficient and powerful tool to bridge the gap between structural biology and system dynamics.

### Anatomy Does Not Equate to Function

A critical insight from our conflict simulation is that anatomical connectivity does not equate to functional efficacy. Although the connectome contains strong inhibitory connections directly to the motor neuron (MN9), our model reveals that these pathways remain functionally silent during escape. Instead, the brain selectively recruits a specific, redundant ensemble to target the premotor source (Roundup). This finding serves as a crucial caution against inferring function solely from synaptic weights without considering dynamic state variables.

### Computational Predictions Guide Experimental Dissection

Finally, this work illustrates the unique value of “virtual genetics.” In biological experiments, identifying redundant mechanisms is notoriously difficult because silencing a single neuron often yields no phenotype, leading to false negatives. Our computational approach enabled us to perform combinatorial perturbations that are technically difficult in vivo, revealing the hidden fault-tolerant architecture of the feeding suppression circuit. These predictions provide precise coordinates for future research: rather than searching for a single “command neuron” that stops feeding, experimentalists should investigate distributed ensembles. By illuminating these invisible architectural principles, computational connectomics provides the necessary blueprint to guide the dissection of complex biological systems.

## Methods

### Connectome Data and Neuron Selection

All neuronal reconstruction and synaptic connectivity data were obtained from the FlyWire full-brain connectome (Codex version 783). Neurons of interest (e.g., LC4, LPLC2, DNp01, MN9) were identified using cell-type annotations in the FlyWire Codex.

### Computational Model

We implemented a Leaky Integrate-and-Fire (LIF) [20, 26] spiking neural network model of the entire central brain, following the framework established by Shiu et al. [20]. The dynamics of the membrane potential *V*_*i*_ for each neuron *i* are governed by the following differential equations:

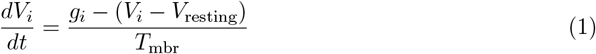

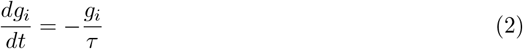

where *g*_*i*_ represents the synaptic conductance resulting from the aggregate firing of presynaptic neurons. Upon the arrival of a spike from presynaptic neuron *j*, the conductance is updated according to the rule: *g*_*i*_ ← *g*_*i*_ + *w*_*j,i*_. When *V*_*i*_ reaches the firing threshold *V*_threshold_, the neuron spikes, and *V*_*i*_ is reset to *V*_reset_ for a refractory period *T*_refractory_.

Biophysical parameters were fixed to values established in previous Drosophila modeling studies [20, 27]:

- Resting potential: *V*_resting_ = −52 mV
- Reset potential: *V*_reset_ = −52 mV
- Firing threshold: *V*_threshold_ = −45 mV
- Membrane time constant: *T*_mbr_ = 20 ms (derived from membrane resistance *R*_mbr_ = 10 kΩ *·* cm^2^ and capacitance *C*_mbr_ = 2*μ*F *·* cm^−2^)
- Synapse decay time constant: *τ* = 5 ms
- Refractory period: *T*_refractory_ = 2.2 ms
- Synaptic delay: *T*_dly_ = 1.8 ms

The synaptic weight *w*_*j,i*_ connecting neuron *j* to neuron *i* is defined as:

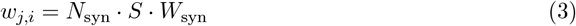

where *N*_syn_ is the number of ultrastructural synapses from *j* to *i* derived from the FlyWire connectome, *S* is the neurotransmitter sign (+1 for excitatory, −1 for inhibitory) predicted by the connectome-wide classifier [23], and *W*_syn_ = 0.275 mV is the global scaling parameter representing the postscynaptic potential change per synapse.

### Simulation strategy

Sensory inputs (Visual looming, Sugar) were modeled as Poisson spike trains delivered to specific receptor neurons (LC4/LPLC2, Sugar GRNs). “Counterfactual” silencing was performed via “virtual axotomy”: the outgoing synaptic weights of specific target neurons (e.g., DNge031) were set to zero, effectively removing their influence on the network while preserving their internal dynamics. All simulations were performed using the Brian2 simulator.

## Data Availability

The neuronal connectivity data used in this study are available from the FlyWire Codex.

## Code Availability

The simulation code and analysis scripts are available at: https://github.com/WJXI/escape-of-Drosophila.

## 1 Acknowledgments

This work was supported by Research Grants Council of Hong Kong (GRF 17311322 and CRF C7012-21GF), National Natural Science Foundation of China (Grant No. 12222416), National Key R&D Program of China (Grants No. 2022YFA1403700), NSFC (Grants No. 12141402) and the Science, Technology and Innovation Commission of Shenzhen Municipality (No. ZDSYS20190902092905285)

